# Education Shapes the Link Between EEG Aperiodic Components and Cognitive Aging

**DOI:** 10.1101/2025.07.02.662700

**Authors:** Sara Lago, Sara Zago, Sonia Montemurro, Rocco Salvatore Calabrò, Maria Grazia Maggio, Serena Dattola, Ilaria Casetta, Giorgio Arcara

**Affiliations:** IRCCS San Camillo Hospital, Venice, Italy; Padova Neuroscience Center, University of Padova, Italy; Department of Philosophy, Sociology, Education and Applied Psychology (FISPPA), University of Padova, Italy; IRCCS Centro Neurolesi “Bonino Pulejo”, Messina, Italy; Department of General Psychology, University of Padova, Italy

**Keywords:** aperiodic components, exponent, offset, healthy aging, cognitive performance, education

## Abstract

Healthy aging brings widespread shifts in aperiodic (non-oscillatory) electroencephalographic (EEG) components, which may underlie physiological changes in cognitive performance. Education, a known protective factor against age-related decline in cognitive performance, has been largely overlooked in studies linking aperiodic EEG components to cognition. This study addresses this gap, hypothesizing that education moderates the interplay between age, aperiodic components, and cognitive performance, as measured by Mini-Mental State Examination (MMSE) scores. We reanalyzed an open-source EEG dataset of 714 healthy individuals aged 18–91 years using Generalized Additive Mixed Models. Aperiodic exponent and offset both declined with age, but higher education levels mitigated these declines. Notably, exponent and offset interacted with age and education in predicting MMSE performance in the bilateral cingulate, left hippocampus, bilateral parietal, right occipital, and left temporal regions. Among older adults, the relationship between the aperiodic components and cognitive performance diverged by education: those with lower education showed worse cognitive outcomes with lower exponents and offsets, whereas higher-educated individuals after 60 years showed a reverse pattern, with lower exponents and offsets predicting better MMSE performance. Our findings suggest that the link between aperiodic components and cognitive aging is not straightforward but depends on moderating factors such as education. These results underscore the importance of accounting for individual differences, like educational background, when exploring age-related changes in EEG aperiodic components and cognition.

## 1. Introduction

Recent research on the EEG signal has shifted from the traditional focus on oscillatory activity in specific EEG frequency bands to a growing interest in the aperiodic (non-oscillatory) component. In the electrophysiological signal, oscillatory activity is associated with peaks that can be observed in the Power Spectrum Density (PSD). However, the PSD is always characterized by a general (and broadband) component, that has been named recently “aperiodic component”, characterized by a 1/f-like distribution. Separating the aperiodic and the periodic component is now a relevant perspective with important implications in the interpretation of results. Historically, neural activity has been measured by averaging power within predefined frequency bands derived from the power spectrum, thus confounding the periodic with the aperiodic component (1). The aperiodic component of the spectrum can be parameterized by the exponent, reflecting the slope of the power spectrum, and the offset, indicating a broadband shift in power across frequencies (1,2). These parameters are associated with two underlying neurophysiological processes: the exponent has been linked with the excitation-to-inhibition balance, reflecting the proportion of excitatory versus inhibitory neural activity in the brain, while the offset has been related to the neural spiking rate, potentially reflecting the overall level of neural activity or baseline firing rates (3,4). However, the relation with a specific underpinning is questioned and several changes in properties (e.g., presence of bursts, or noise in the signal) can affect the aperiodic component of the spectrum (5–7).

Recent studies have demonstrated the functional significance of aperiodic components, associating variations in its parameters with task performance (3,8,9), age-related cognitive decline (10–13), and brain disorders (14,15). Consequently, the aperiodic component is increasingly recognized as a critical biomarker for understanding neural processes and cognitive function (14,16).

Aging is associated with widespread changes in aperiodic EEG components that reflect non-pathological alterations in neuronal networks (10,12). Both the exponent and offset of the aperiodic component tend to decline with age, indicating shifts in the excitation-to-inhibition (E:I) balance and reductions in overall neural firing rates, respectively (10,12,17,18).

Age-related changes in aperiodic components contribute to cognitive performance throughout the lifespan. Specifically, the reduction in the exponent (often referred to as the flattening of the aperiodic slope) has been linked to age-related declines in cognitive functions, such as working memory (12). Subsequent studies have replicated these findings and linked flatter slopes in older adults to poorer performance on spatial attention, short-term memory, processing speed, and executive function tasks(9,11,19,20). Additionally, beyond the exponent, the offset has been found to be negatively correlated with reaction time, perceptual sensitivity, processing speed, and selective attention performance (8,20–22). These findings support the neural noise hypothesis, which posits that aging-related reductions in the exponent reflect increased desynchronized background neural activity, or neural noise, driven by shifts in the E:I ratio. This increased noise disrupts neural communication fidelity, leading to a flatter power spectrum and reduced efficiency in cognitive processing (12). Consequently, decreases in the exponent during healthy aging may indicate greater neural excitability and diminished E:I balance, aligning with broader patterns of age-related neurophysiological change (23).

However, some findings contradict these results. Euler et al. (21) reported no association between the aperiodic exponent and specific cognitive constructs such as working memory, perceptual reasoning, processing speed, and verbal comprehension in participants aged 18 to 52. However, they observed a relationship between the exponent and an aggregated measure of cognitive ability. Similarly, Cesnaite et al. (24) found no link between aperiodic components and cognitive performance in an older cohort (aged 60–80), while Smith et al. (25) reported that higher exponents were associated with better general cognitive function but were unrelated to age (50–80).

That said, these associations are not consistent across all ages and tasks, suggesting that the relationship between aperiodic components and cognition is task- and age-dependent (24,26) and highlighting the possible influence of unexamined moderating variables such as education. Education is a well-established predictor of cognitive performance, with higher educational attainment typically associated with better cognitive outcomes (27,28). While some studies found no moderating effect of education on the relationship between aperiodic components and cognition (24), preliminary findings suggest otherwise. Montemurro et al. (13) reported that in older adults, the relationship between the aperiodic exponent and cognitive performance varied by education: higher exponents were predictive of worse processing speed and working memory in highly educated individuals, but better performance in those with lower education. The authors suggest that the relationship between exponent, neural noise, and cognitive performance may not be straightforward, and highlight the importance of investigating possible mediators, such as education, within this complex relationship. However, this study categorized participants into discrete groups (e.g., younger vs. older adults, higher vs. lower education), treating inherently continuous variables as categorical. This approach reduces data variability and may obscure nuanced relationships between education, aging, and aperiodic components. To address such existing limitations and further explore the potential moderating role of education in the relationship between aperiodic components and cognition in aging, we reanalyzed data from a publicly available dataset (29) (accessible at https://github.com/euroladbrainlat/Brain-health-in-diverse-setting). This dataset includes measures of aperiodic EEG components (exponent and offset), age, education, and Mini-Mental State Examination (MMSE) scores (30) as continuous variables for 714 participants aged 18 to 91 years, thus covering a wider age range than previous studies (21,24,25).

The present study aimed to achieve two objectives: [1] to examine the relationships between EEG aperiodic components, age, and education, with the goal of confirming established age-related changes and investigating less-studied effects of education on aperiodic components, and [2] to investigate whether demographic variables (age and education) and EEG aperiodic components could predict cognitive performance measured with the MMSE. We hypothesized that education may serve as a protective factor against age-related decline in aperiodic exponent and offset. Furthermore, we posited that this protective effect might extend to cognitive abilities, such that participants with higher educational attainment would exhibit higher aperiodic exponents and offsets, alongside better MMSE performance, compared to their less-educated counterparts.

## 2. Materials and Methods

### 2.1. Participants

Demographic and cognitive data for the sample are summarized in Table 1. From the original larger dataset, a subsample of 714 participants for which MMSE and aperiodic EEG data were available were included in our analyses. Participants had no history of psychiatric or neurological disorders, alcohol or drug abuse, significant sensory impairments, or functional cognitive complaints. Additional details about participants are provided in Hernandez et al. (29).

**Table 1.**
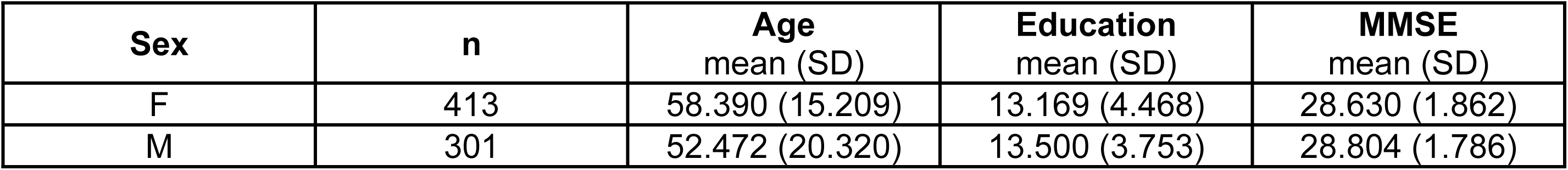
Demographics and cognitive information for the participants’ sample.

| Sex | n | Age<br>mean (SD) | Education<br>mean (SD) | MMSE<br>mean (SD) |
| --- | --- | --- | --- | --- |
| F | 413 | 58.390 (15.209) | 13.169 (4.468) | 28.630 (1.862) |
| M | 301 | 52.472 (20.320) | 13.500 (3.753) | 28.804 (1.786) |

### 2.2. EEG Acquisition and Processing

Details of EEG acquisition, preprocessing, source estimation, and aperiodic component extraction are described in Hernandez et al. (29). Briefly, after recording the EEG at rest (both in eyes-closed and eyes-open conditions) and preprocessing, EEG data was filtered between 0.5 and 40 Hz. Source reconstruction was performed using the standardized low resolution electromagnetic tomography (sLORETA) algorithm (31), projecting the EEG signal on the Montreal Neurological Institute (MNI) template. The source-reconstructed signal was segmented into 82 regions using the Automated Anatomical Labeling (AAL) atlas (32).

The aperiodic exponent and offset were derived from the source-reconstructed EEG signal using the FOOOF algorithm, which models the aperiodic component with a Lorentzian function:

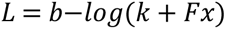

where *b* represents the broadband offset, *χ* is the exponent, and *k* is the ‘knee’ parameter, which accounts for the bend in the aperiodic component (not considered in the present analyses). F denotes the vector of input frequencies (1).

To facilitate data interpretation and enhance the identification of patterns in the results, these regions were aggregated into ten composite regions of interest (ROIs) via mean averaging. The composite ROIs and their corresponding anatomical regions are presented in Table 2. The dataset did not include information on the aperiodic components in the frontal brain areas.

**Table 2.**
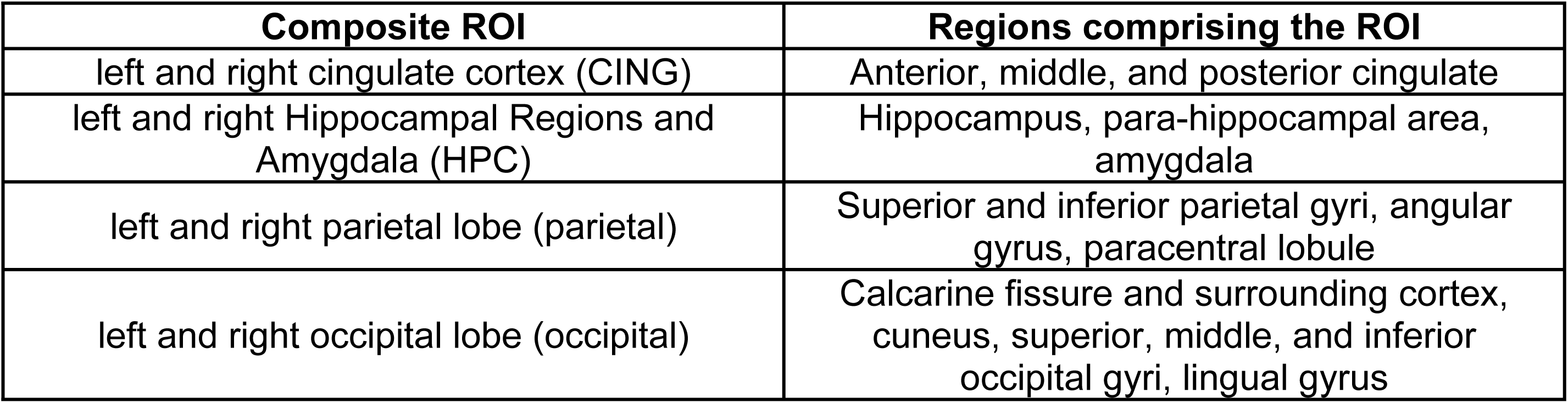
Corresponding brain regions for each composite ROI (adapted from Hernandez et al. (29)).

| Composite ROI | Regions comprising the ROI |
| --- | --- |
| left and right cingulate cortex (CING) | Anterior, middle, and posterior cingulate |
| left and right Hippocampal Regions and Amygdala (HPC) | Hippocampus, para-hippocampal area, amygdala |
| left and right parietal lobe (parietal) | Superior and inferior parietal gyri, angular gyrus, paracentral lobule |
| left and right occipital lobe (occipital) | Calcarine fissure and surrounding cortex, cuneus, superior, middle, and inferior occipital gyri, lingual gyrus |

### 2.3 Statistical analyses

Statistical analyses were conducted using R (version 4.4.1; 14.06.2024;(33)). Generalized Additive Mixed Models (GAMMs) were implemented using the mgcv package (version 1.9.1; (34)). GAMMs are nonlinear mixed-effects regression methods (35,36) and were chosen because they allow to model nonlinear interactions between variables. Moreover, they can also accommodate continuous variables (37), a crucial feature when investigating cognitive and brain aging such in the present case: Thuwal et al. (19) pointed out that changes in exponent are associated in a non-linear way with age, and aging is indeed a continuous process, that accumulates effects gradually, and can assume different individual trajectories (38). Therefore, these features need to be considered in the statistical model.

Separate models were computed for each region of interest (ROI) and each aperiodic EEG component (exponent and offset). To account for individual differences, all models included a random factor smooth for subjects, enabling nonlinear adjustments to regression shapes (36). The negative binomial family was used in all models to address non-normality in residuals. For both research aims, a model comparison strategy was employed to identify the most appropriate statistical model for each ROI. Separate simple and complex models were constructed for each aim and ROI.

For Aim 1 (exploring relationships between EEG components, age, and education) the following models were constructed:

● Simple: EEG component ∼ sex + s(Age) + s(Education) + s(ID, bs=’re’)
● Complex: EEG component ∼ sex + s(Age) + s(Education) + ti(Age, Education) + s(ID, bs=’re’)

The simple model evaluated the main effects of age and education, while the complex model additionally considered their interaction.

On the other hand, for Aim 2 (exploring how changes in MMSE scores relate to EEG components, age, and education) the following models were constructed:

● Simple: transformed MMSE scores ∼ sex + s(EEG component) + s(Age) + s(Education) + s(ID, bs=’re’)
● Complex: transformed MMSE scores ∼ sex + s(EEG component) + s(Age) + s(Education) + ti(EEG component, Age) + ti(EEG component, Education) + ti(Age, Education) + ti(EEG component, Age, Education) + s(ID, bs=’re’)

In Aim 2, the simple model did not include any EEG components and was therefore independent of specific ROIs. This model captured known patterns of cognitive change across increasing age and education levels and served as a "baseline" against which all other complex models were compared.

In all models, the number of basis functions (the k parameter) was set to the default values: 9 for main effects, 16 for two-way interactions, and 64 for three-way interactions. These values provided sufficient model flexibility given the available data while preventing overfitting (see the Deviance Explained columns of Tables 3-6).

**Table 3.**
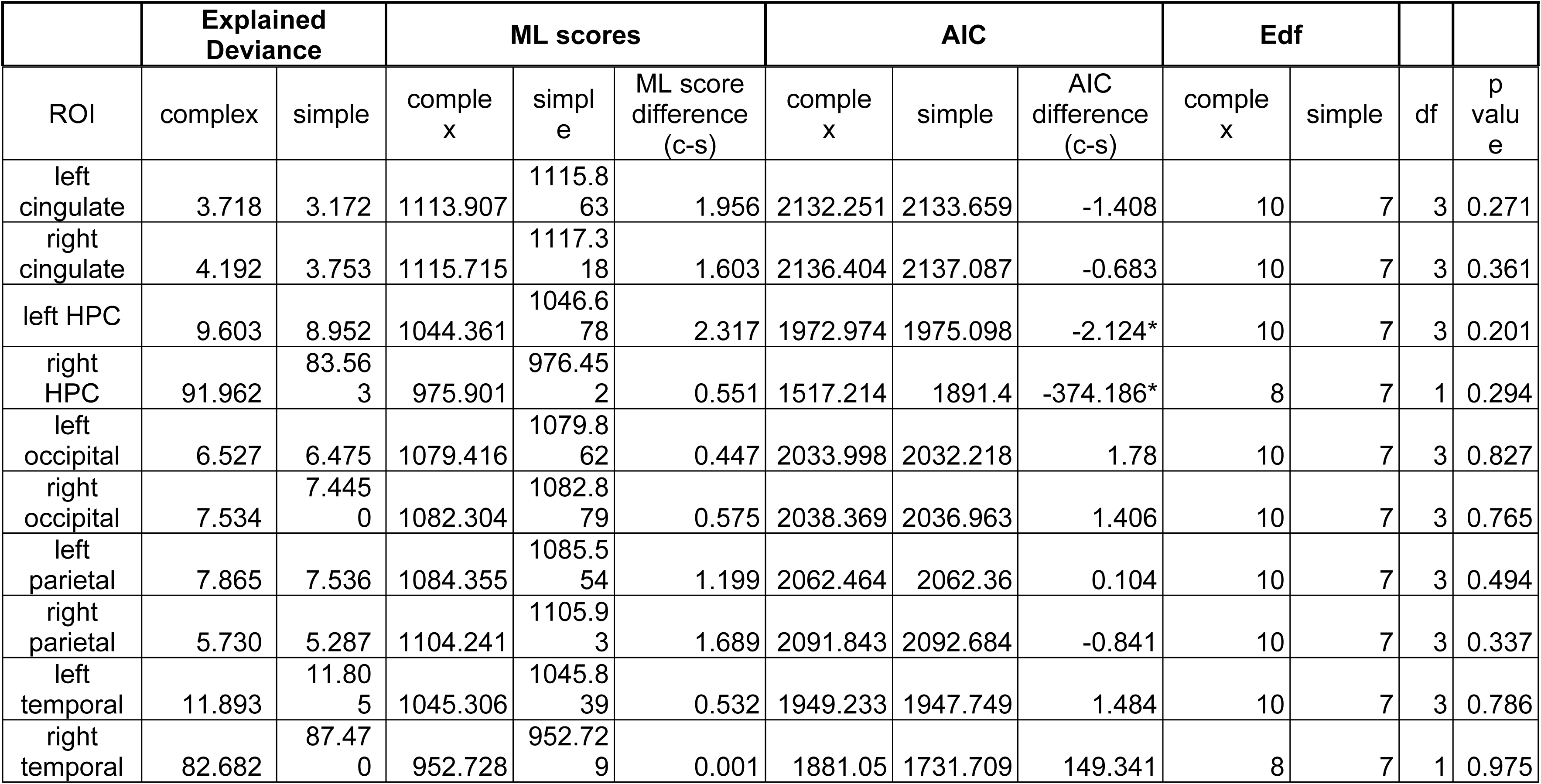
ML and AIC scores (including differences between complex and simple models), along with effective degrees of freedom (Edf) and p-values assessing the significance of differences between the simple and complex exponent models. Percentages of explained deviance are also reported. Model selection is only based on AIC differences greater than -2 (marked with *).

| ROI | complex | simple | complex | simple | ML score difference (c-s) | complex | simple | AIC difference (c-s) | complex | simple | df | p value |
| --- | --- | --- | --- | --- | --- | --- | --- | --- | --- | --- | --- | --- |
| left cingulate | 3.718 | 3.172 | 1113.907 | 1115.863 | 1.956 | 2132.251 | 2133.659 | -1.408 | 10 | 7 | 3 | 0.271 |
| right cingulate | 4.192 | 3.753 | 1115.715 | 1117.318 | 1.603 | 2136.404 | 2137.087 | -0.683 | 10 | 7 | 3 | 0.361 |
| left HPC | 9.603 | 8.952 | 1044.361 | 1046.678 | 2.317 | 1972.974 | 1975.098 | -2.124* | 10 | 7 | 3 | 0.201 |
| right HPC | 91.962 | 83.563 | 975.901 | 976.452 | 0.551 | 1517.214 | 1891.4 | -374.186* | 8 | 7 | 1 | 0.294 |
| left occipital | 6.527 | 6.475 | 1079.416 | 1079.862 | 0.447 | 2033.998 | 2032.218 | 1.78 | 10 | 7 | 3 | 0.827 |
| right occipital | 7.534 | 7.4450 | 1082.304 | 1082.879 | 0.575 | 2038.369 | 2036.963 | 1.406 | 10 | 7 | 3 | 0.765 |
| left parietal | 7.865 | 7.536 | 1084.355 | 1085.554 | 1.199 | 2062.464 | 2062.36 | 0.104 | 10 | 7 | 3 | 0.494 |
| right parietal | 5.730 | 5.287 | 1104.241 | 1105.93 | 1.689 | 2091.843 | 2092.684 | -0.841 | 10 | 7 | 3 | 0.337 |
| left temporal | 11.893 | 11.805 | 1045.306 | 1045.839 | 0.532 | 1949.233 | 1947.749 | 1.484 | 10 | 7 | 3 | 0.786 |
| right temporal | 82.682 | 87.470 | 952.728 | 952.729 | 0.001 | 1881.05 | 1731.709 | 149.341 | 8 | 7 | 1 | 0.975 |

It is worth noticing that MMSE scores exhibit a strong left-skew due to the high prevalence of scores near the upper limit (30) among healthy participants, particularly younger individuals. Including such a skewed variable in the models caused significant deviations from normality in the models’ residuals, which compromised the reliability of inferences derived from the results. To address this issue and bring the residuals closer to normality, MMSE scores were reversed and log-transformed prior to their inclusion in the model.

Model comparisons were conducted using the Akaike Information Criterion (AIC) (39): difference of more than -2 indicated a better fit for the more complex model. Only the results from the best-fitting complex models—those with an AIC difference greater than -2—are reported here, while results for simpler models are available in the Supporting Information. For completeness, Maximum Likelihood (ML) scores, model comparisons based on ML, and their corresponding p-values are also reported alongside AIC values for each model. However, model selection was based solely on AIC differences.

#### 2.3.1 Significance Testing for predicted effects

Significance testing followed the approach of Sóskuthy (40). This approach is mainly based on the visual interpretation of each model’s main effects or interaction surfaces and the relative surface differences, while the main effects’ and interactions’ (also called smoothers) p-values are not considered. After identifying the best-fitting model based on AIC, for simple models significant main effects were assessed by plotting smoothers (available in the Supporting Information) with a customized version of the plot_smooth function. For Aim1, these plots illustrated aperiodic components depending on age or education as continuous variables (terms s(Age) and s(Education) in the models); for Aim 2, they illustrated MMSE scores depending on aperiodic components (term s(EEG component)), age or education. In both cases, they resulted in 2D graphs.

For complex models instead, the interactions’ plots resulted in 3D graphs (surface plots). For Aim 1, they depicted aperiodic components depending on the two-way association between age and education (term ti(Age, Education), Figures 1 and 3), while for Aim 2 they depicted the MMSE scores depending on three-way association between aperiodic components, age and education (term ti(EEG component, Age, Education), Figures 5a and 6). Surface plots were produced using a customized version of vis.gam. In the surface plots, lighter yellow and green shades correspond to higher values of the response variable (exponent, offset or MMSE score), while darker blue shades correspond to lower values of the response variable. White areas indicate regions where the confidence intervals (95%) around the predicted surface included zero, i.e., where the interaction was not significant.

**Figure 1.**
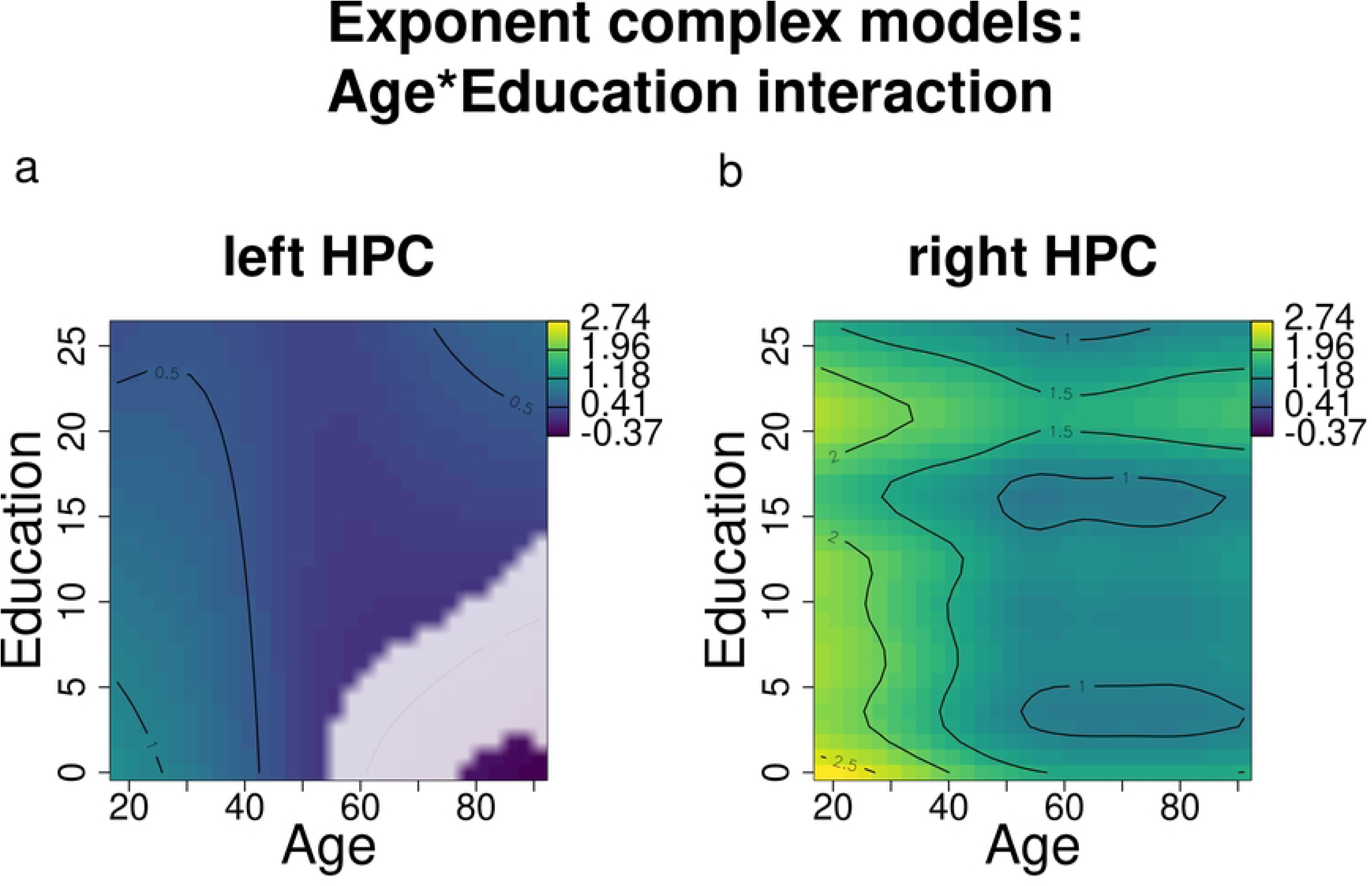
Exponent changes over age and education (results of the complex models’ interactions) for the left and right HPC. Darker blue shades indicate lower exponents, while lighter green and yellow shades indicate higher exponents. White areas indicate regions where the confidence intervals (95%) around the predicted surface included zero, i.e., the interaction was not significant.

Since the interpretation of such 3D graphs can be confusing, it is possible to break them down by selecting different values of education (for Aim 1 plots) or aperiodic components (for Aim 2 plots). This allows us to plot 2D graphs “zooming in” on the dependent variable (aperiodic component or MMSE score, as per the different aims) in relation with selected levels of an independent variable: we can examine how exponent and offset change according to different education levels with increasing age (Aim 1, Figures 2a-b, 4a-d) and how MMSE scores change according to different education and exponent levels with increasing age (Aim 2, Figures 7a-d). These plots were obtained with the plot_smooth function.

**Figure 2.**
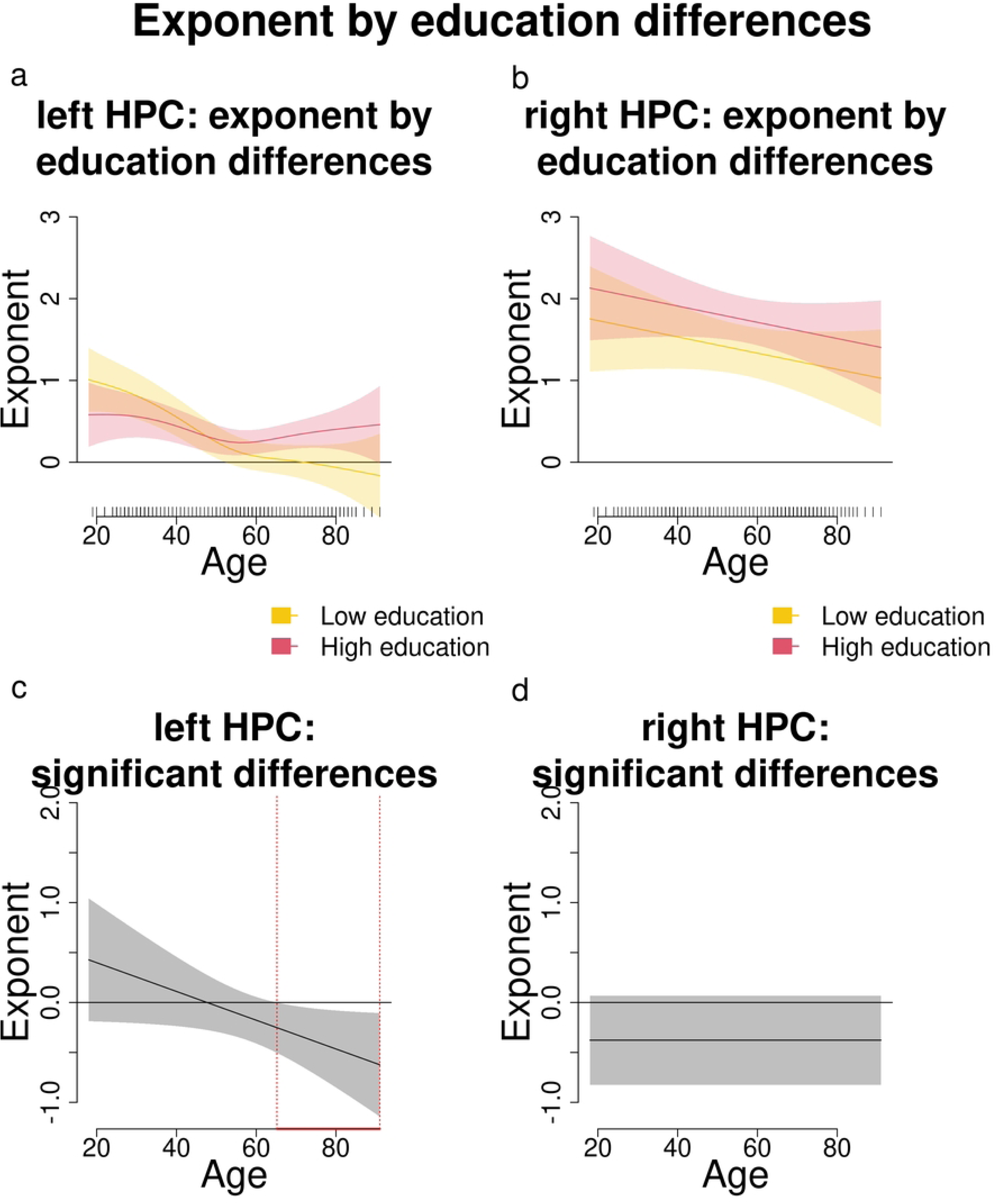
Comparisons and significant differences between exponent levels of participants with high and low education, in relation with age for the left and right HPC. Panel a and b show how age shapes the exponent for different levels of education (high and low). Panel c and d show significant differences in exponent levels between participants with varying levels of education across age. Age ranges when differences are significant are highlighted with a red line on the x axis. Panels a and c show that in the left HPC participants with high education have significantly higher exponents than those with lower education starting approximately from age 65. No significant differences are reported in the right HPC (panels b and d).

From these 2D representations we can compute significant differences in the dependent variable according to the different selected levels of the independent variable. In our case, for Aim 1 we can examine whether there is a significant effect of education on exponent and offset, and at which age ranges this difference is significant (Figures 2c-d, 4e-h; 4m-p); in the same vein, for Aim 2 we can test whether and at which age range the interaction between education and exponent has a significant effect on MMSE scores (Figures 7e-h). In these difference plots, age ranges of significant differences are highlighted with a red line on the x axis. These plots were obtained using the plot_diff function. [40]. The R functions for plotting and significance testing are available in the package itsadug (version 2.4.1; (41)).

## 3. Results

### 3.1. Aim 1: relationships between aperiodic components, age and education

#### 3.1.1. Exponent

When analyzing the exponent, a complex model incorporating the interaction term was found to be preferable only for the bilateral HPC. In other regions, either the differences in AIC between models were negligible or the inclusion of the interaction term negatively impacted the model’s fit (e.g., right temporal; Table 3).

Age and education modulated the exponent differently in the bilateral HPC, as illustrated in Figure 1, where higher exponents are represented by lighter yellow and green shades, while lower exponents are represented by darker blue shades. Younger individuals generally exhibited higher exponents, which declined with age (in Figure 1a-b, the lighter shades in correspondence to young age become darker with increasing age). Importantly, education appeared to mitigate these age-related decreases (in the upper right corner of Figure 1a, darker blue in correspondence to older age and lower education becomes lighter with increasing education; in Figure 1b, blue becomes green/yellow in correspondence with education levels around 20 years).

Comparisons between individuals with high and low education levels (Figure 2) revealed that differences in exponent became significant only in the left HPC after the age of 65 (Figure 2c). This suggests that in the left HPC, education significantly modulates the relationship between age and exponent, with higher education levels associated with less pronounced declines in exponent values over time.

In summary, we found that older adults exhibit lower exponents (flatter slopes) in the left and right HPC. Furthermore, older adults with higher education demonstrated significantly higher exponents in the left HPC compared to their peers with lower education levels.

#### 3.1.2. Offset

When analyzing the offset, a complex model incorporating the interaction between age and education was found to be preferable over a simpler model in several regions, including the right cingulate, bilateral HPC, left occipital, and bilateral parietal and temporal regions. However, in the left cingulate, adding the interaction term increased the AIC, indicating a poorer fit, while in the right occipital, the interaction term led to only a negligible improvement in AIC (Table 4).

**Table 4.** ML and AIC scores (including differences between complex and simple models), along with effective degrees of freedom (Edf) and p-values assessing the significance of differences between the simple and complex offset models. Percentages of explained deviance are also reported. Model selection is only based on AIC differences greater than -2 (marked with *).

| ROI | Explained Deviance |  | ML scores |  |  | AIC |  |  | Edf |  | df | p value |
| --- | --- | --- | --- | --- | --- | --- | --- | --- | --- | --- | --- | --- |
|  | complex | simple | complex | simple | ML score difference (c-s) | complex | simple | AIC difference (c-s) | complex | simple |  |  |
| left cingulate | 37.620 | 37.857 | 1436.165 | 1439.505 | 3.34 | 3028.754 | 3015.259 | 13.495 | 10 | 7 | 3 | 0.083 |
| right cingulate | 81.062 | 76.769 | 1426.044 | 1429.797 | 3.752 | 2823.097 | 2908.163 | -85.066* | 10 | 7 | 3 | 0.057 |
| left HPC | 94.195 | 84.050 | 1468.387 | 1472.689 | 4.302 | 2284.215 | 2851.016 | -566.801* | 10 | 7 | 3 | 0.035 |
| right HPC | 88.918 | 75.724 | 1456.577 | 1460.264 | 3.688 | 2640.162 | 3000.336 | -360.174* | 10 | 7 | 3 | 0.061 |
| left occipital | 65.023 | 64.864 | 1411.902 | 1412.038 | -0.136 | 2867.159 | 2947.831 | -80.672* | 7 | 10 | 3 | 1.000 |
| right occipital | 41.704 | 42.119 | 1423.33 | 1423.65 | 0.32 | 3002.261 | 3003.272 | -1.011 | 10 | 7 | 3 | 0.887 |
| left parietal | 88.404 | 88.262 | 1359.893 | 1360.846 | 0.953 | 2456.79 | 2463.91 | -7.12* | 10 | 7 | 3 | 0.592 |
| right parietal | 77.851 | 77.176 | 1411.435 | 1412.946 | 1.511 | 2783.5 | 2834.343 | -50.843* | 10 | 7 | 3 | 0.388 |
| left temporal | 78.817 | 72.006 | 1443.121 | 1444.784 | 1.663 | 2932.754 | 3023.847 | -91.093* | 10 | 7 | 3 | 0.344 |
| right temporal | 82.681 | 81.234 | 1440.388 | 1441.863 | 1.474 | 2839.875 | 2874.256 | -34.381* | 10 | 7 | 3 | 0.4 |

Age and education were observed to interact in modulating offset levels differently across regions (Figure 3). In the left occipital and bilateral parietal regions, variations in offset across different age and education levels were minimal, with values constrained within a narrow range (between 1 and 2; Figures 3d-f). Conversely, in regions such as the right cingulate, bilateral HPC, and bilateral temporal areas, younger age and higher education were associated with increased offset levels, mirroring trends observed for the exponent (Figure 3a-b,g-h).

**Figure 3.**
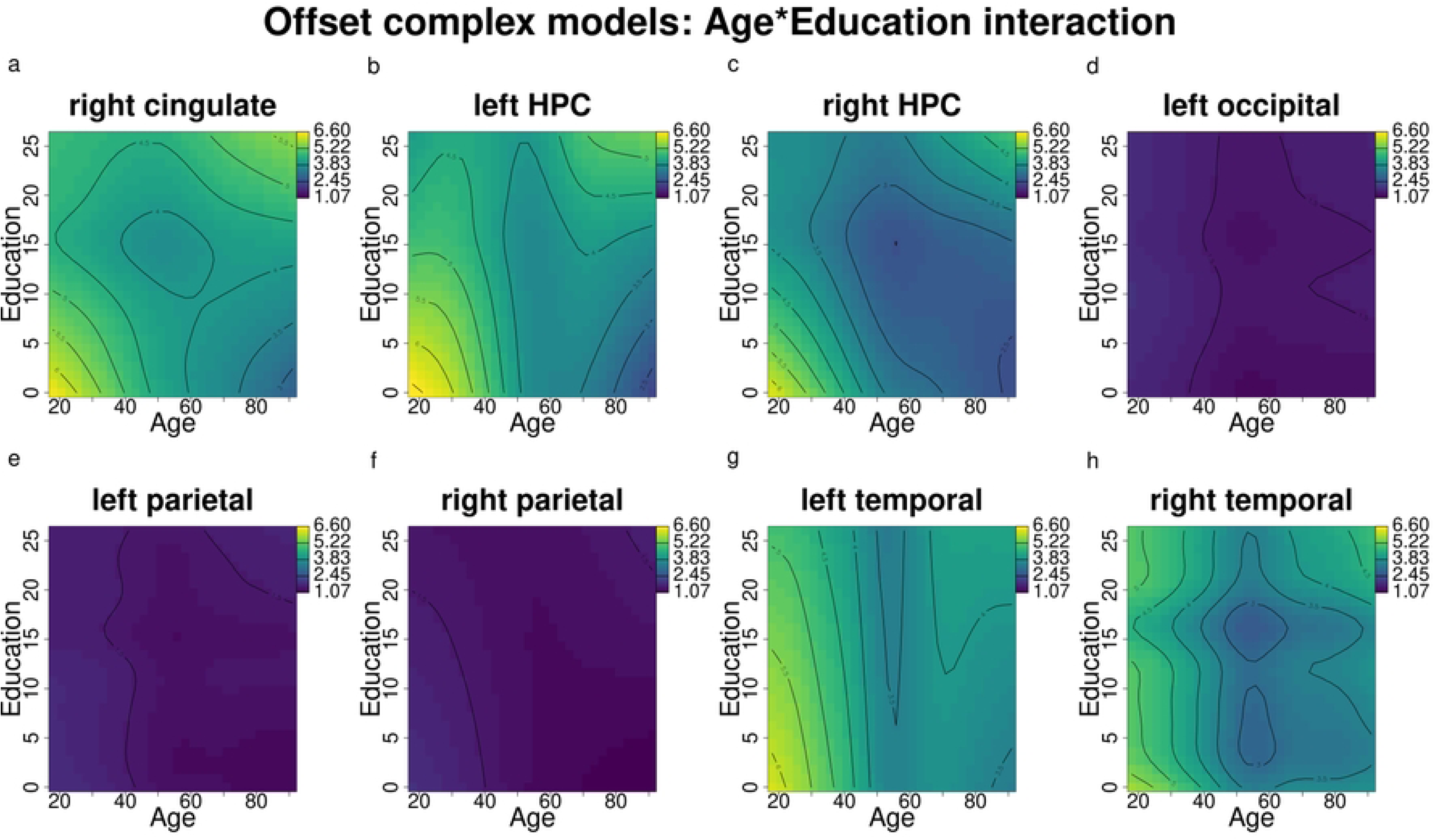
Offset changes according to age and education (results of the complex models’ interactions) for the right cingulate, left and right HPC, left occipital, left and right parietal, and left and right temporal. Darker blue shades indicate lower offsets, while lighter green and yellow shades indicate higher offsets.

Direct comparisons between individuals with higher and lower education levels revealed that older individuals with higher education exhibited significantly higher offset levels compared to their peers with lower education levels in the right cingulate, bilateral HPC, and bilateral parietal regions, where a small but significant difference can be observed (Figure 4a–c, 4i-j). Differences were significant within the age range of 65 to 91 years.

**Figure 4.**
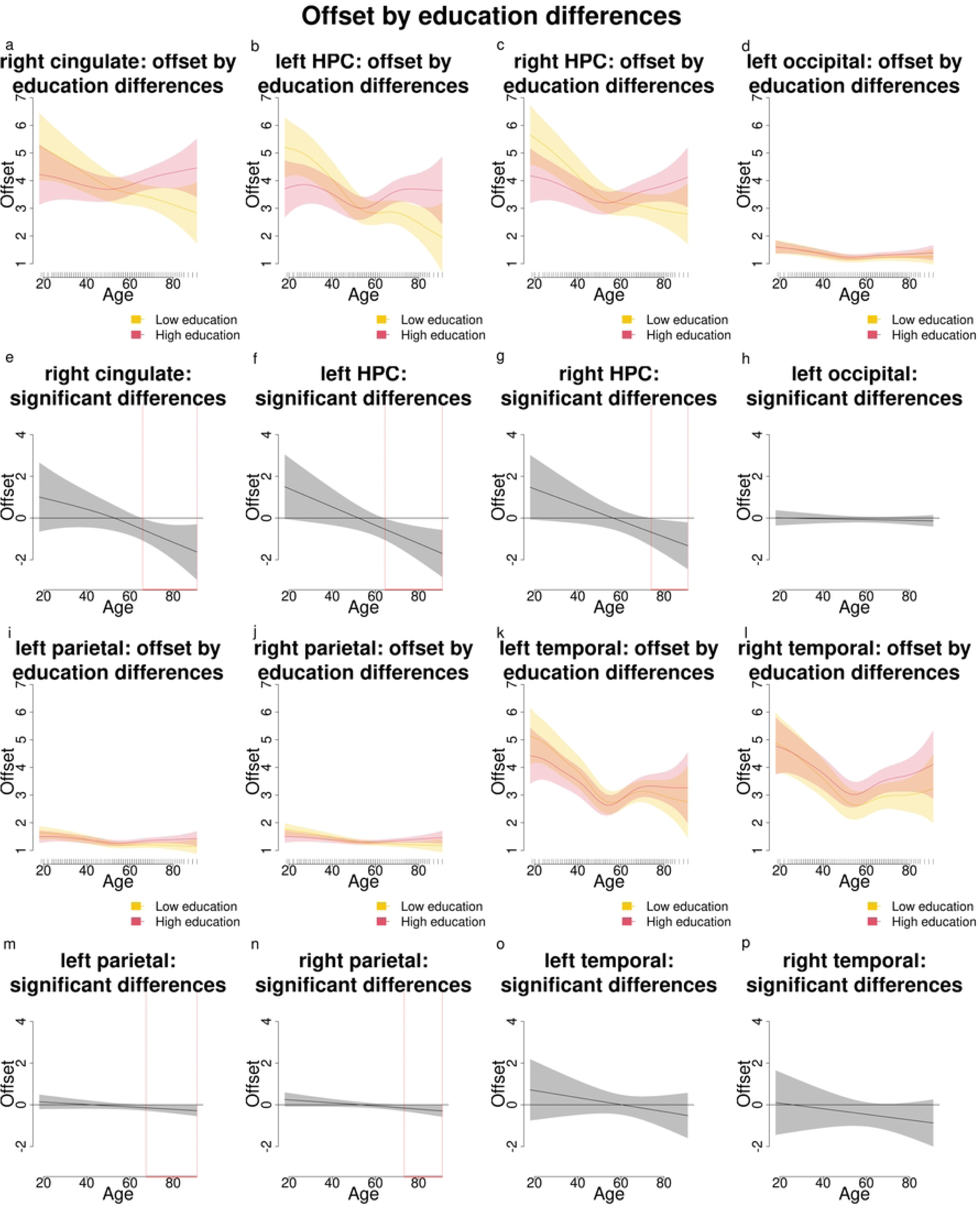
Comparisons and significant differences between offset levels of participants with high and low education, in relation with age for the right cingulate, bilateral HPC, left occipital, bilateral parietal, and bilateral temporal areas. Panels a-d, i-l show how age shapes the offset for different levels of education (high and low). Panels e-h, m-p show significant differences in offset between participants with varying levels of education across age. Age ranges when differences are significant are highlighted with a red line on the x axis. Panels a-c, i and j show that in the right cingulate, bilateral HPC and bilateral parietal participants with high education have significantly higher offsets than those with lower education starting approximately from age 65 or older. No significant differences are reported in the left occipital and bilateral parietal areas (panels d,h,o,p).

In summary, in regions such as the right cingulate, bilateral HPC and bilateral parietal, education was found to significantly modulate offset levels in a manner comparable to its influence on exponent levels observed in the bilateral HPC.

#### 3.1.3. Aim 1 results summary

Analyses aiming at exploring relationships between EEG components, age, and education revealed that in the left HPC, older adults exhibited lower exponents (flatter slopes), but higher education mitigated the age-related decrease in exponent values. A similar pattern was observed for the offset in regions such as the right cingulate, bilateral HPC, and bilateral parietal areas.

### 3.2. Aim 2: aperiodic components, age, and education as predictors of MMSE scores

#### 3.2.1. Baseline MMSE model

The baseline MMSE model revealed a significant interaction between age and education in influencing MMSE scores. As expected, younger participants and older participants with higher education had higher MMSE scores compared to older participants with lower education (yellow and green shades as opposed to darker blue shades in Figure 5a; see Figure 5b for direct comparisons between participants with higher and lower education). The difference between participants with higher and lower education was significant across the entire age range of interest (20 to 91 years old; Figure 5c).

**Figure 5.**
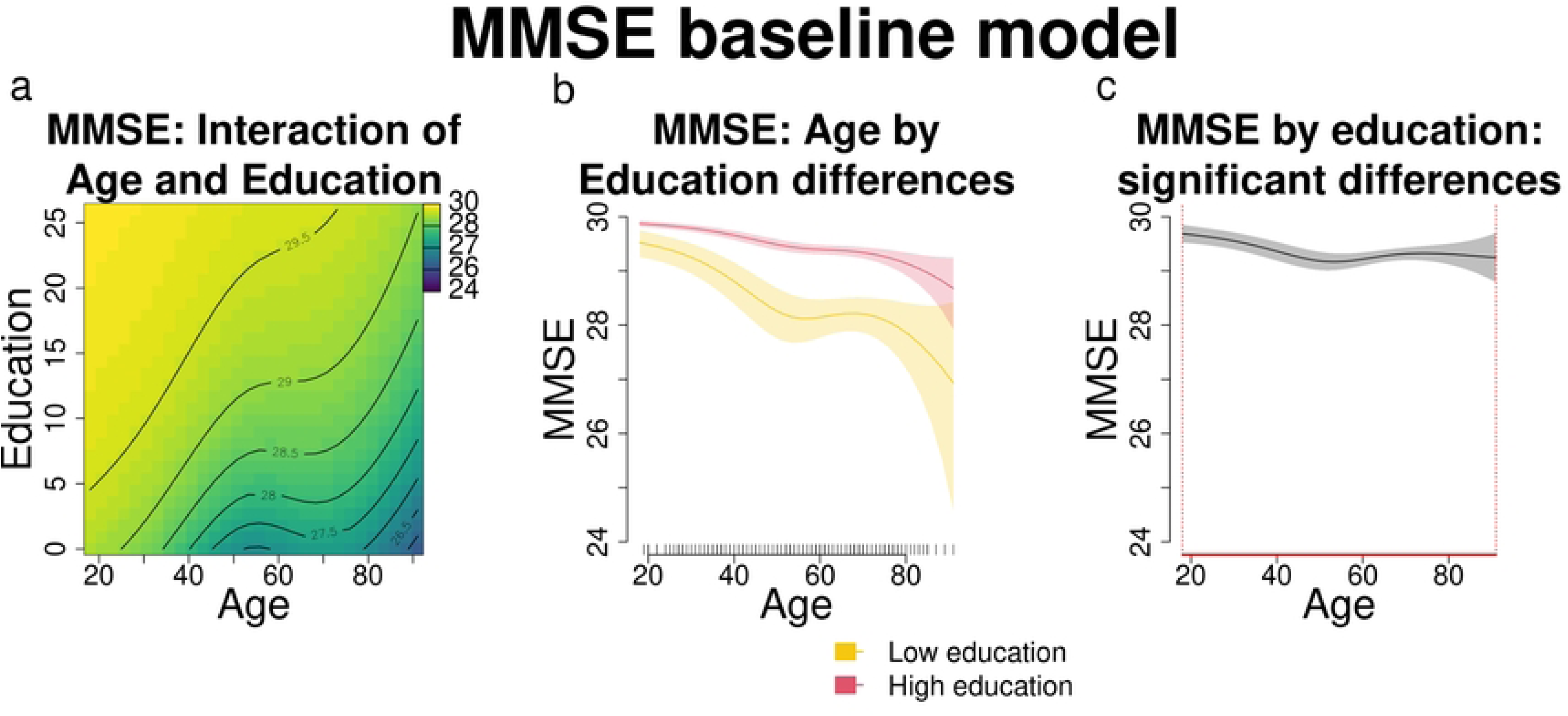
a) MMSE scores changes according to age and education (results of the models’ interaction). Darker blue shades indicate lower MMSE scores, while lighter green and yellow shades indicate higher scores. b) Comparison between MMSE scores of participants with high and low education, in relation with age: varying levels of education shape MMSE performance differently across age. c) Significant differences between MMSE scores of participants with high and low education, in relation with age. Age ranges when differences are significant are highlighted with a red line on the x axis.

#### 3.2.2. MMSE and exponent

When examining the relationship between MMSE scores and exponent, a more complex model incorporating the interaction term between age, education, and exponent was preferred over the baseline MMSE model in the bilateral cingulate, left hippocampal, right occipital, bilateral parietal and left temporal regions (Table 5). In all these ROIs, the exponent influences the known interaction between age, education, and MMSE scores: for lower exponents (Figure 6a,d), the interaction effect mirrors that of the baseline MMSE model (Figure 5a), where younger age and older age with higher education are associated with higher MMSE scores. Moving towards median exponent values (Figure 6b,e), higher MMSE scores are observed even among older individuals with lower education; however, at higher exponent levels (Figure 6c,f), this association appears to reverse: whereas older participants with higher education had higher MMSE scores at lower and median exponent levels, older individuals with higher exponents and higher education show worse MMSE scores compared to those with lower education. Since the effects are relatively consistent across all regions, only the figures for the left and right cingulate are presented here, while figures for the remaining regions can be found in the Supporting Information.

**Figure 6.**
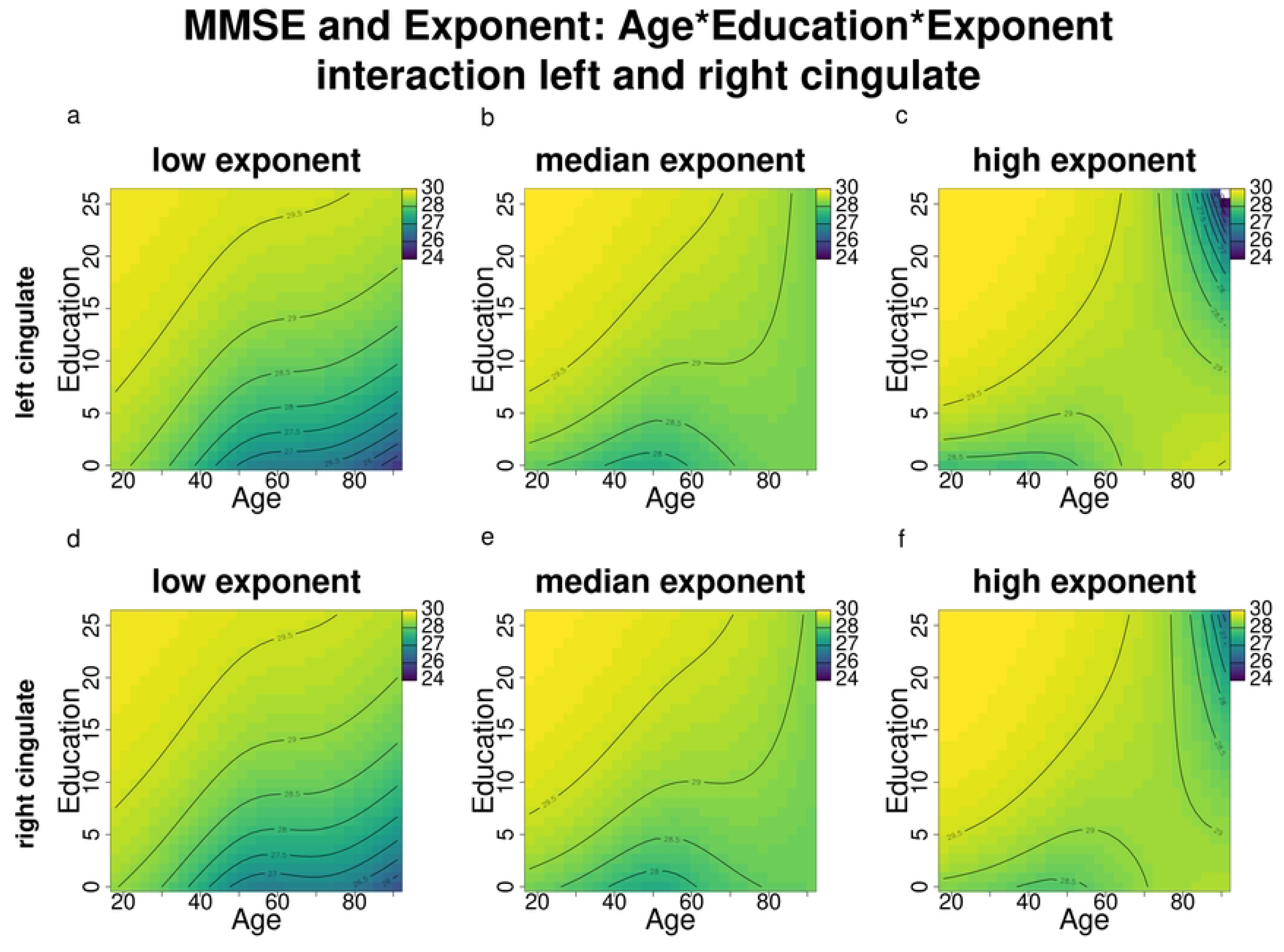
MMSE score changes according to age and education across different exponent levels (results of the complex models’ interactions) for the bilateral cingulate. Darker blue shades indicate lower MMSE scores, while lighter green and yellow shades indicate higher MMSE scores.

**Table 5.** ML and AIC scores (including differences between complex and simple models), along with effective degrees of freedom (Edf) and p-values assessing the significance of differences between the baseline MMSE model and complex models incorporating the interaction with the exponent. Percentages of explained deviance are also reported. Model selection is only based on AIC differences greater than -2 (marked with *).

| ROI | Explained Deviance |  | ML scores |  |  | AIC |  |  | Edf |  | df | p value |
| --- | --- | --- | --- | --- | --- | --- | --- | --- | --- | --- | --- | --- |
|  | complex | simple | complex | simple | ML score difference (c-s) | complex | simple | AIC difference (c-s) | complex | simple |  |  |
| left cingulate | 13.388 | 11.233 | 921.266 | 928.903 | 7.637 | 1269.955 | 1275.768 | -5.813* | 19 | 10 | 9 | 0.084 |
| right cingulate | 13.289 | 11.233 | 922.004 | 928.903 | 6.899 | 1270.902 | 1275.768 | -4.866* | 19 | 10 | 9 | 0.13 |
| left HPC | 12.486 | 11.233 | 922.866 | 928.903 | 6.899 | 1273.298 | 1275.768 | -2.47* | 19 | 10 | 9 | 0.209 |
| right HPC | 12.055 | 11.233 | 924.502 | 928.903 | 6.899 | 1275.459 | 1275.768 | -0.309 | 19 | 10 | 9 | 0.456 |
| left occipital | 12.612 | 11.233 | 923.329 | 928.903 | 5.574 | 1274.649 | 1275.768 | -1.119 | 19 | 10 | 9 | 0.266 |
| right occipital | 12.930 | 11.233 | 922.816 | 928.903 | 6.087 | 1272.769 | 1275.768 | -2.999* | 19 | 10 | 9 | 0.204 |
| left parietal | 12.947 | 11.233 | 922.076 | 928.903 | 6.827 | 1272.281 | 1275.768 | -3.487* | 19 | 10 | 9 | 0.135 |
| right parietal | 12.710 | 11.233 | 922.649 | 928.903 | 6.254 | 1272.823 | 1275.768 | -2.945* | 19 | 10 | 9 | 0.186 |
| left temporal | 13.001 | 11.233 | 923.356 | 928.903 | 5.547 | 1272.912 | 1275.768 | -2.856* | 19 | 10 | 9 | 0.269 |
| right temporal | 12.107 | 11.233 | 924.22 | 928.903 | 4.683 | 1274.982 | 1275.768 | -0.786 | 19 | 10 | 9 | 0.404 |

When comparing individuals with high and low exponents across different education levels, this reversed association becomes clearer (Figure 7). Among participants with lower education, those with higher exponents have higher MMSE scores than those with lower exponents, particularly at middle and older ages (approximately >40; Figure 7a,c). In contrast, among participants with higher education, this pattern is reversed: participants with higher exponents show higher MMSE scores at younger ages (approximately <60), but lower MMSE scores at older ages (Figure 7b,d). These differences are significant across the entire age range of interest (18-91 years; Figure 7e-h). Another important observation is the greater variability in MMSE scores among participants with lower education compared to those with higher education. In contrast, their counterparts with higher education, MMSE scores are clustered near the upper limit (30) until approximately age 65, after which scores of participants with higher exponents begin to decline.

**Figure 7.**
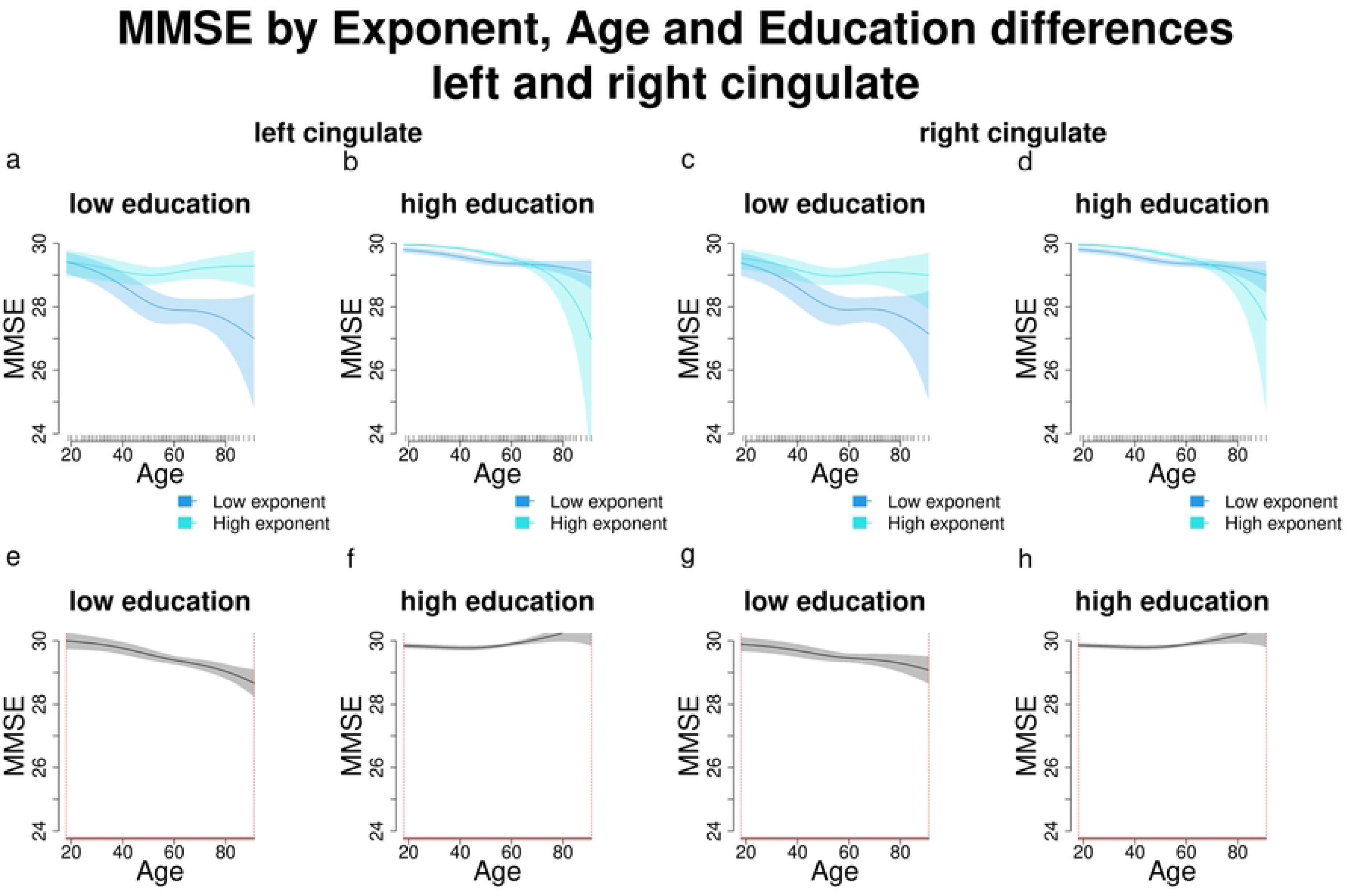
Comparisons and significant differences between MMSE scores of participants with high and low exponents, in relation with age and different education levels for the left and right cingulate. Panels a-d show how different exponent levels shape MMSE scores across different levels of education (high and low) and increasing age. Panels e-h show significant differences in MMSE scores between participants with varying exponent and education levels across age. Age ranges when differences are significant are highlighted with a red line on the x axis. Panels a and c show that in the left and right cingulate, participants with low education and higher exponents have significantly higher MMSE scores than those with lower exponents starting approximately from age 40. Panels b and d show that participants with higher education and higher exponents have worse MMSE scores than those with lower exponents. In panels e-h age ranges when differences are significant are highlighted with a red line on the x axis (MMSE score differences are significant across all the age window).

In summary, in older adults with lower education, there is a possible positive relationship between exponent and MMSE scores (lower exponents correspond to lower scores), while those with higher education show the opposite trend (lower exponents correspond to higher MMSE scores).

#### 3.2.3. MMSE and offset

When examining the relationship between MMSE scores and offset, the pattern of results closely mirrors the findings on MMSE and exponent. More complex models that included the interaction between age, education, and exponent were preferred over the baseline MMSE model in the bilateral cingulate, left hippocampus, right occipital, bilateral parietal, and left temporal regions (Table 6).

**Table 6.** ML and AIC scores (including differences between complex and simple models), along with effective degrees of freedom (Edf) and p-values assessing the significance of differences between the baseline MMSE model and complex models incorporating the interaction with the offset. Percentages of explained deviance are also reported. Model selection is only based on AIC differences greater than -2 (marked with *).

| ROI | Explained Deviance |  | ML scores |  |  | AIC |  |  | Edf |  | df | p value |
| --- | --- | --- | --- | --- | --- | --- | --- | --- | --- | --- | --- | --- |
|  | complex | simple | complex | simple | ML score difference (c-s) | complex | simple | AIC difference (c-s) | complex | simple |  |  |
| left cingulate | 13.329 | 11.233 | 920.79 | 928.903 | 8.113 | 1269.41 | 1275.768 | -6.358* | 19 | 10 | 9 | 0.062 |
| right cingulate | 13.459 | 11.233 | 921.201 | 928.903 | 7.702 | 1269.937 | 1275.768 | -5.831* | 19 | 10 | 9 | 0.08 |
| left HPC | 12.997 | 11.233 | 921.287 | 928.903 | 7.616 | 1269.69 | 1275.768 | -6.078* | 19 | 10 | 9 | 0.085 |
| right HPC | 12.278 | 11.233 | 923.856 | 928.903 | 5.047 | 1275.208 | 1275.768 | -0.56 | 19 | 10 | 9 | 0.343 |
| left occipital | 12.4196 | 11.233 | 923.283 | 928.903 | 5.62 | 1274.482 | 1275.768 | -1.286 | 19 | 10 | 9 | 0.26 |
| right occipital | 12.757 | 11.233 | 922.987 | 928.903 | 5.916 | 1273.636 | 1275.768 | -2.132* | 19 | 10 | 9 | 0.223 |
| left parietal | 13.152 | 11.233 | 921.362 | 928.903 | 7.541 | 1271.322 | 1275.768 | -4.446* | 19 | 10 | 9 | 0.089 |
| right parietal | 12.665 | 11.233 | 922.451 | 928.903 | 6.452 | 1273.085 | 1275.768 | -2.683* | 19 | 10 | 9 | 0.167 |
| left temporal | 13.012 | 11.233 | 923.038 | 928.903 | 5.865 | 1272.78 | 1275.768 | -2.988* | 19 | 10 | 9 | 0.229 |
| right temporal | 11.951 | 11.233 | 924.326 | 928.903 | 4.577 | 1275.785 | 1275.768 | 0.017 | 19 | 10 | 9 | 0.423 |

As in the previous section, since the effects are relatively consistent across all regions, we report only the figures for the left and right cingulate here, while figures for the remaining regions are available in the Supporting Information. Offset influences the established relationship between age, education, and MMSE scores in a manner similar to its effect on the relationship between MMSE and exponent. Specifically, at lower offset values (Figure 8a,d), the interaction effect resembles that of the baseline MMSE model (Figure 5a). As offset values increase toward the median (Figure 8b,e), higher MMSE scores are observed even among older individuals with lower education. However, at higher offset levels (Figure 8c,f), this association reverses: older individuals with both higher offsets and higher education exhibit lower MMSE scores compared to those with lower education.

**Figure 8.**
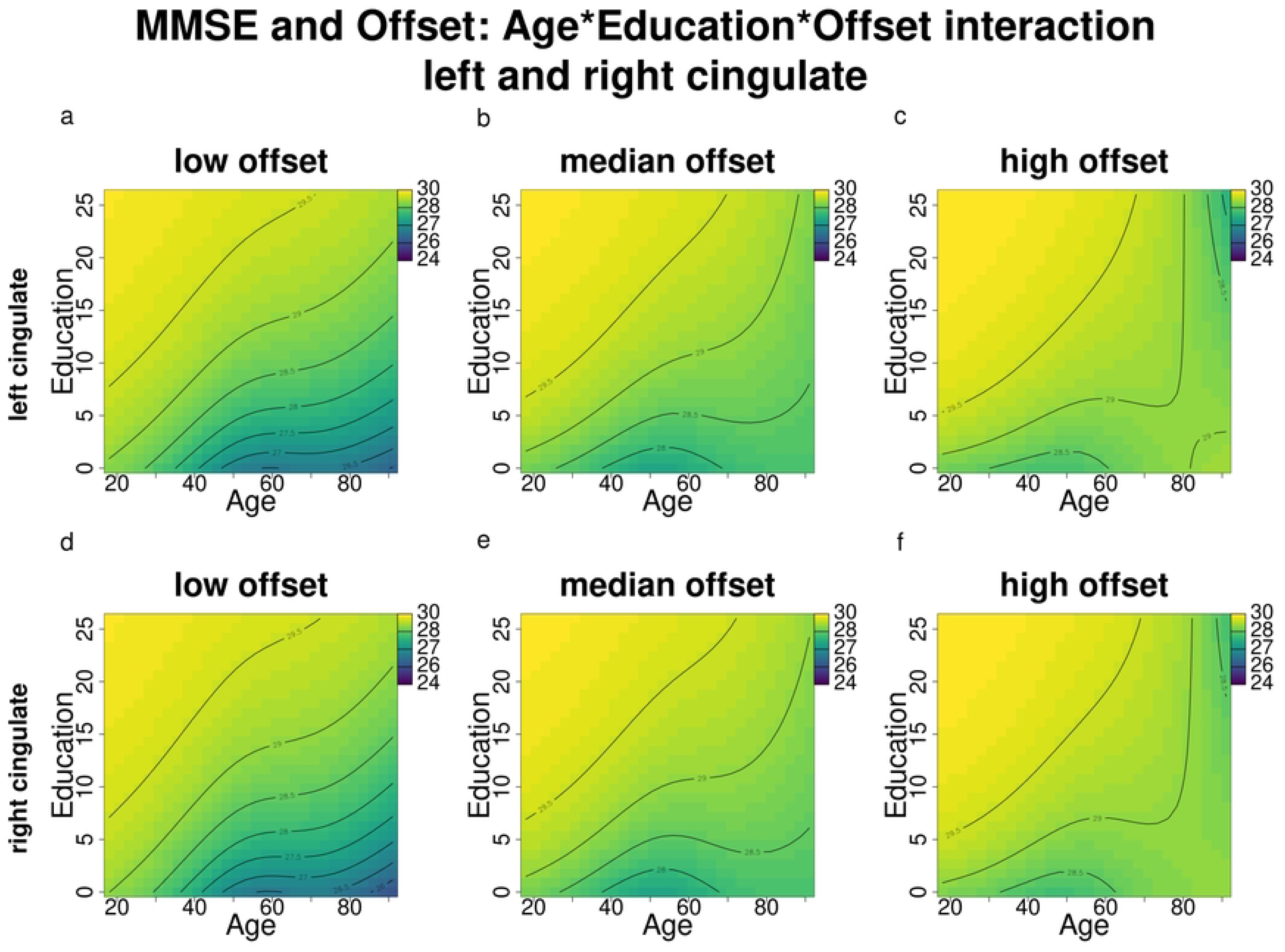
MMSE score changes according to age and education across different offset levels (results of the complex models’ interactions) for the bilateral cingulate. Darker blue shades indicate lower MMSE scores, while lighter green and yellow shades indicate higher MMSE scores.

When comparing participants with high and low offsets across different education levels (Figure 9), we observe distinct trends. Among participants with lower education, those with higher offsets tend to show better MMSE scores beginning around age 40 (Figure 9a,c). However, this pattern reverses in participants with higher education: individuals with higher offsets display higher MMSE scores until approximately age 60, after which their scores decline (Figure 9b,d). These differences are statistically significant across the entire age range of interest (18–91 years; Figure 9e-h) and closely parallel the relationship between MMSE and exponent across varying levels of age and education.

**Figure 9.**
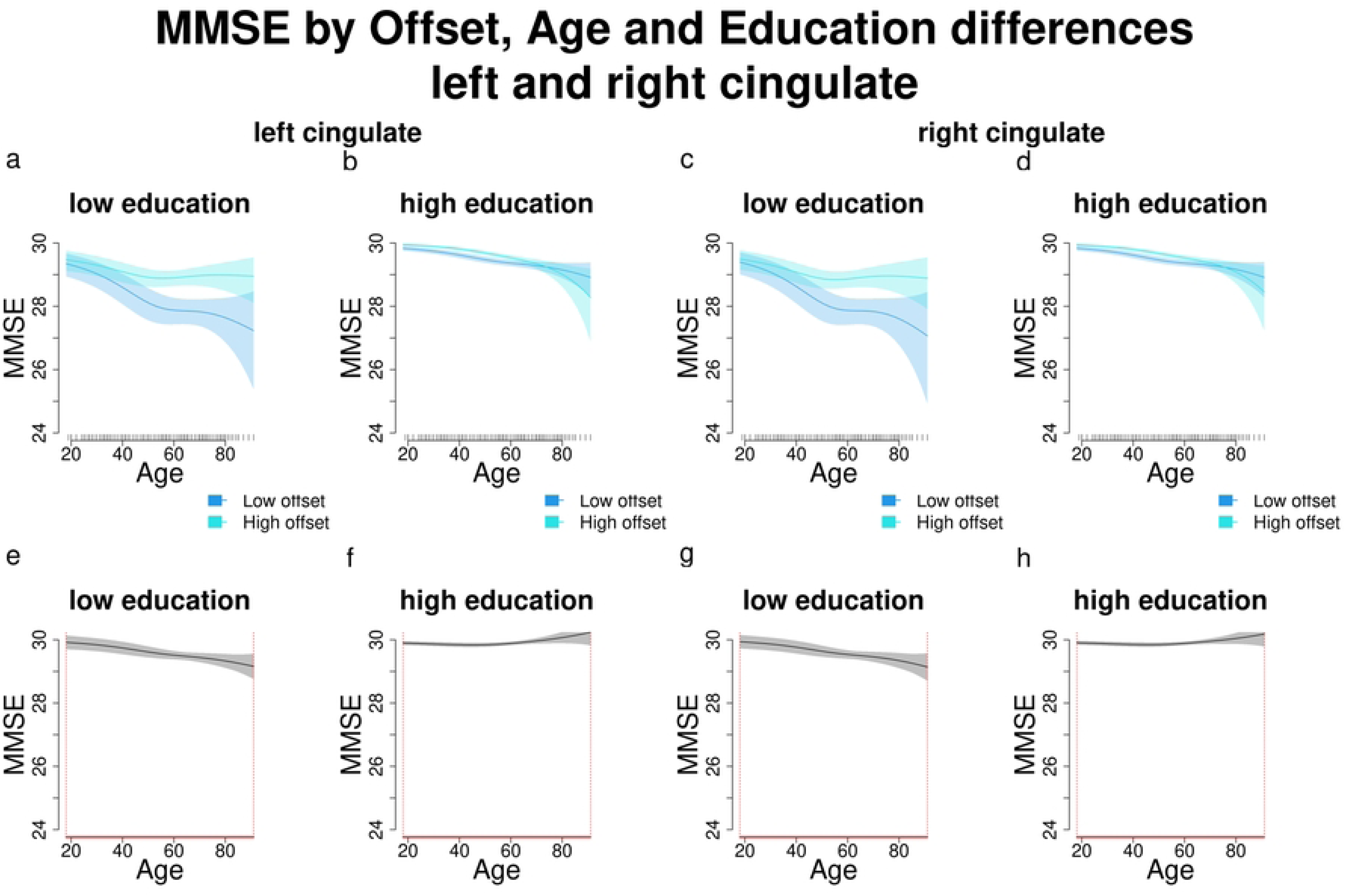
Comparisons and significant differences between MMSE scores of participants with high and low offsets, in relation with age and different education levels for the left and right cingulate. Panels a-d show how different offset levels shape MMSE scores across different levels of education (high and low) and increasing age. Panels e-h show significant differences in MMSE scores between participants with varying offset and education levels across age. Age ranges when differences are significant are highlighted with a red line on the x axis. Panels a and c show that in the left and right cingulate, participants with low education and higher offsets have significantly higher MMSE scores than those with lower offsets starting approximately from age 40. Panels b and d show that participants with high education and higher exponents have worse MMSE scores than those with lower exponents. In panels e-h age ranges when differences are significant are highlighted with a red line on the x axis (MMSE score differences are significant across all the age window).

#### 3.2.4. Aim 2 results summary

The results for the aperiodic exponent and offset in relation to MMSE scores were remarkably similar: in both cases, more complex models that included interaction terms between the aperiodic component, age, and education were preferred over the baseline MMSE model in the bilateral cingulate, left hippocampus, bilateral parietal, right occipital, and left temporal regions. Additionally, the pattern of results remained consistent across both ROIs and aperiodic components.

More in detail, in the above-mentioned areas both the exponent and offset significantly predicted cognitive performance across age and education levels. In older adults with lower education, a positive relationship between the aperiodic components and MMSE scores was observed, with lower exponents and offsets corresponding to lower MMSE scores. In contrast, among those with higher education, a reverse trend was found, where lower exponents and offsets were associated with higher MMSE scores.

## 4. Discussion

The present study addressed two main objectives: [1] to examine the relationships between EEG aperiodic components, age, and education, aiming to confirm known age-related changes while exploring the less-studied effects of education on aperiodic components; and [2] to investigate whether age, education, and EEG aperiodic components could predict cognitive performance, as measured by MMSE scores.

Our findings align with the existing literature on age-related reductions in the exponent and offset, and on the association between reductions in the aperiodic components and worsening cognition (5,6,8,9,16–19). However, they also revealed that age and education interacted in predicting the exponent in the bilateral hippocampal areas and the offset in multiple brain regions, including the right cingulate, bilateral hippocampus, bilateral occipital, parietal, and temporal regions, extending the findings of previous studies (8,9,12,20,22). Notably, we found that individuals with higher educational attainment exhibited higher exponents and offsets than their less-educated peers between the ages of approximately 65 and 91, suggesting a possible protective effect of education against age-related declines in the exponent and offset.

Our findings also add other novel insights: both the exponent and offset interacted with age and education in predicting MMSE scores in the bilateral cingulate, left hippocampus, bilateral parietal, right occipital, and left temporal regions. Participants with lower education showed a positive relationship between both aperiodic components and MMSE performance, with lower exponents and offsets predicting worse cognitive outcomes as age increased. Conversely, participants with higher education displayed the opposite trend starting at around age 60, where lower aperiodic components were associated with better MMSE performance. This pattern of results should be considered with caution due to the low inter-individual variability in MMSE scores among highly educated younger and middle-aged participants, whose scores clustered near the upper limit (30). In contrast, older participants and those with lower education exhibited greater variability in cognitive performance. This limited variability in MMSE scores among highly educated individuals may partially explain the observed differences in cognitive performance between participants with high and low education (see Figures 5b and 7a-d). Despite this limitation, our findings suggest that higher education likely modulates the association between aperiodic components and MMSE performance, and that these aspects could jointly act as protective factors against cognitive decline in older age. Previous studies investigating the relationship between education and aperiodic components (21,26) found no significant effects, even when using large datasets, including the one reanalyzed here (26). Those studies primarily used linear regression and cluster-based correlation methods, which may not have captured the non-linear relationships instead identified by our approach. Our results instead align with and extend the findings by Montemurro et al. (13), despite methodological differences. While they grouped participants by age and education and employed linear regression, we treated age and education as continuous variables and used a flexible statistical model. Consistent with their findings, we observed that older adults with lower education levels benefited cognitively from higher exponents and offsets, while higher-educated older adults showed the opposite pattern.

Strictly regarding the MMSE performance across age (baseline MMSE model results), our results reaffirm the well-established link between higher education and better cognitive performance (Figure 5). According to a threshold model of cognitive decline, individuals with higher levels of education exhibit better cognitive functioning compared to those with lower education throughout the entire age span and are able to compensate for age-related cognitive declines over a longer period (their cognitive performance drops later in life) (27). The cognitive reserve theory extends this view, adding that highly educated individuals’ might exhibit slower rates of cognitive decline (28,42). Figures 5b, 7b, and 7d show that higher-educated participants’ MMSE scores clustered near the upper limit (30), while those with lower education exhibited greater variability, supporting the observation that on average, individuals with higher education also have a better MMSE performance throughout the entire age span. From age 60 onwards instead, the decline in MMSE scores was steeper in participants with lower education, consistent with the cognitive reserve framework, suggesting a causal role of education (related to further life experiences, e.g., complexity of the occupational level) that may influence the capacity of brain structure and functions to cope with normal and pathological age-related changes (28,42).

Adding the aperiodic components to the baseline MMSE model (Section 3.2.2. and 3.2.3.; MMSE and exponent, MMSE and offset) reversed the association between education and cognitive performance in older age (Figure 7b,d). Among older adults with lower education, higher exponents and offsets predicted better cognitive performance and less pronounced cognitive decline. In contrast, higher-educated older adults exhibited the reverse pattern: higher aperiodic components predicted worse performance and steeper decline, as observed by Montemurro et al. (13) for the exponent. These findings also align with the neural noise hypothesis, which posits that reduced exponents reflect increased neural noise due to an increased E:I ratio during healthy aging (12). Lower exponents are thus associated with flatter power spectra, decreased neural communication fidelity, and greater excitability (higher noise). These associations between aperiodic exponent and E:I ratio are supported by in-silico models (43) and studies manipulating the neural noise (measured with the aperiodic exponent) through the administration of pharmacological agents that either decrease (e.g., propofol) or increase excitation (ketamine) (20,43,44). Age-related changes in the offset also fit the neural noise hypothesis in that the offset is functionally related to the underlying neural population’s spiking rate (3,4); diminishing offsets and, therefore, diminishing synchronous spiking rates could partly explain the age differences in aperiodic components observed here and in a growing body of research (12,13,18,19,45).

Interestingly, the stochastic resonance framework (46) suggests that there may be no single "optimal" level of neural noise for cognitive performance. Instead, the effects of noise on cognition may depend on individual factors and compensatory dynamics, so that subjective changes in the aperiodic exponent, and thus in noise levels and E:I balance, might perturbate the cognitive system’s functioning in an age- and education-dependent way, according to the individual’s unique characteristics (47). Further research is needed to clarify the role of aperiodic offset in age-related changes in neural noise levels. While its exact influence remains uncertain, it is possible that age-related alterations in synchronous neural spiking contribute to the modulation of neural noise (4,12,18).

The neural noise and stochastic resonance hypotheses may therefore complement the cognitive reserve theory in explaining how the interaction between the aperiodic components and education influences cognitive performance, particularly in older adults. Our findings show that in older individuals with lower education, lower exponents and offsets are associated with poorer MMSE performance. This is consistent with previous research (9,11,12,19,20) and parallels findings in younger participants, where lower exponents are linked to poorer performance on complex tasks (11,48). This pattern likely reflects the physiological decline associated with aging. In contrast, highly educated older adults maintain preserved cognitive performance despite lower exponent and offset values. This suggests that individuals with greater cognitive reserve are better equipped to compensate for both normal and pathological age-related changes at the cognitive and neural level (28,42). In this context, lower aperiodic components—interpreted as increased neural noise and decreased synchronous neural spiking—may reflect compensatory neural processes that are accessible to individuals with higher cognitive reserve but not to those with lower cognitive reserve. In other words, greater neural noise in highly educated individuals may indicate adaptive mechanisms that help sustain cognitive function, supporting the idea that higher educational attainment serves as a protective factor against cognitive and neural decline in older age.

These considerations underscore that the relationship between aperiodic components, age, and cognition is not straightforward, but is mediated by individual traits (47) and especially by education, an important factor in influencing the resilience of the cognitive system to age-related changes. This nonlinearity justifies our use of a statistical approach capable of capturing complex interactions, emphasizing the need for considering individual differences with appropriate analytical tools when studying continuous age-related brain and cognitive changes (38).

It is worth to point out that exponent and offset are highly correlated due to their shared influence on the power spectrum (45,49,50): any exponent change consists in a rotation of the PSD, therefore it might also be related to changes in the offset, even though distinct neural processes likely underlie each of these components (18,50). However, there is an intrinsic ambiguity in the literature regarding the relative role of the offset versus the exponent in the relationship between the aperiodic component and cognitive performance, as well as regarding the expected topographies of the associations between cognitive performance and aperiodic features (21). Indeed, the dataset we use encompasses a broad age range (18–91 years), spanning the entirety of adulthood, which enabled us to reliably investigate the neurodevelopmental dynamics of aperiodic components and their influence on cognitive performance from early adulthood to late life. However, while the temporal patterns provide valuable insights, the spatial distribution of our findings merits further discussion. Age and education interacted in predicting the aperiodic components only in the hippocampal regions and the offset across widespread areas; on the other hand, MMSE scores were significantly associated with these factors across the cortex.

At the present stage, we can only offer a tentative interpretation of this pattern, because the exact neurobiological basis of aperiodic components still remains unclear (21,23): the majority of studies cited in the present work either focused on electrode clusters (10,11,23,48) or averaged all scalp electrodes (21,22,26,45,49), while some focused a-priori on the occipital region (12,13) and others (conducted using magnetoencephalography) found that cognitive performance was associated with aperiodic components in a distributed way across the brain (51). In addition, the study providing the currently used dataset (29) investigated aperiodic components on source-reconstructed EEG data, but aggregated smaller brain areas into composite ROIs (Table 2). Such aggregation reduces the spatial resolution and therefore poses some limitations to the interpretations of our results; another limitation regarding the spatial distribution of the effects is that the currently used dataset did not provide any information about aperiodic exponent and offset within the frontal areas. However, the main theoretical explanations of the functional role of aperiodic components, like the neural noise hypothesis (12), constitute frameworks of general, widespread neural mechanisms that are likely not limited to single brain areas. Further research is therefore needed to clearly establish whether the broad regions in which we found a significant association between MMSE score, aperiodic components and demographic variables are truly more sensitive to age-related aperiodic changes: this caution is necessary in that aging is associated with a widespread increase of low frequencies power, that can result in exponent decrease across the whole brain (7). In addition, EEG source reconstruction algorithms may be inaccurate in estimating activation of mesial areas (52).

Another potential limitation is the use of the MMSE to assess cognitive performance, particularly in younger participants and those with higher education. For these individuals, the MMSE is relatively easy to complete, often resulting in a ceiling effect, where scores cluster near the upper limit (30), thereby reducing inter-individual variability. However, the MMSE was chosen because it remains a widely used cognitive screening tool in both clinical and research settings. Its broad availability, ease of administration, and brevity make it a practical option for initial cognitive assessment, especially in contexts requiring quick and accessible evaluations (53).

Future research should investigate individual differences in cognition using more comprehensive measures that assess a broader range of cognitive domains. A final limitation that it is important to consider is the correlational nature of the study that prevents precise interpretations on the direction of the effect. Although we can hypothesize that education shapes the relationship between aperiodic components and cognition, we cannot exclude the role of other unknown variables that moderate this relationship. This is an intrinsic limitation of cross-sectional studies on aging, that it is worth to underline.

In light of these considerations, we can hypothesize that education might modulate the impact of aging on aperiodic exponent in specific brain regions through mechanisms of cognitive reserve and neuroplasticity. Regions like the hippocampus and cingulate cortex, which are key to memory, attention, and monitoring processes (54,55), may exhibit varying degrees of vulnerability or resilience to age-related changes. This could explain the interaction between age and education in shaping aperiodic components in these areas, with higher education potentially helping to buffer against cognitive decline and preserving neural efficiency in these critical regions. In addition, the fact that the interaction between age and education in predicting the offset was extended to broader regions may rely on the fact that the offset is a measure associated to the frequency of neural firing and it is influenced by global changes in connectivity and synaptic efficiency (4,56); the widespread effect may reflect the idea that education not only acts locally but also promotes greater resilience in broader brain networks. This could result in modulation of the offset across various cortical regions, supporting enhanced neural communication despite aging. This merits further investigation, as well as the neurobiological basis of aperiodic components in general and the interplay between aperiodic components and demographic variables in shaping cognitive performance in specific brain areas.

In conclusion, our findings suggest that the relationship between aperiodic components and cognitive aging is complex and influenced by educational attainment, which may act as a protective factor against age-related decline, especially in the bilateral cingulate, left hippocampus, bilateral parietal, right occipital, and left temporal regions. These results align with the neural noise and stochastic resonance hypotheses (9,46), emphasizing the importance of accounting for individual differences (47) when studying age-related changes in EEG aperiodic components and cognition. Future research should further explore the neurobiological basis of aperiodic components and investigate the reason why specific brain regions seem most sensitive to these interactions.

## Supporting Information captions

The following supporting information can be downloaded at: https://osf.io/x8dnp/?view_only=0c2060b3853540c9a9d1360d38f7d2af, Table S1: Significance of the smooth terms (interactions and main effects of age and education) for the exponent models; Table S2: Significance of the smooth terms (interactions and main effects of age and education) for the offset models; Table S3: Significance of the smooth terms (interactions and main effects of age and education) for the baseline MMSE model; Table S4: Significance of the smooth terms (interactions and main effects of age, education, and exponent) for the MMSE and exponent models; Table S5: Significance of the smooth terms (interactions and main effects of age, education, and offset) for the MMSE and offset models; Figure S1: Main effect of age on exponent for the simple models of left and right cingulate, left and right occipital, left and right parietal, left and right temporal; Figure S2: Main effect of education on exponent for the simple models of left and right cingulate, left and right occipital, left and right parietal, left and right temporal; Figure S3: Main effect of age on offset for the simple models of left and right cingulate; Figure S4: Main effect of education on offset for the simple models of left and right cingulate; Figure S4: MMSE score changes according to age and education across different exponent levels for the left hippocampus; Figure S5: MMSE score changes according to age and education across different exponent levels for the left temporal region; Figure S6: MMSE score changes according to age and education across different exponent levels for the right occipital region; Figure S7: MMSE score changes according to age and education across different exponent levels for the bilateral parietal regions; Figure S8: Comparisons and significant differences between MMSE scores of participants with high and low exponent, in relation with age and different education levels for the left hippocampus; Figure S9: Comparisons and significant differences between MMSE scores of participants with high and low exponent, in relation with age and different education levels for the left temporal region; Figure S10: Comparisons and significant differences between MMSE scores of participants with high and low exponent, in relation with age and different education levels for the left occipital region; Figure S11: Comparisons and significant differences between MMSE scores of participants with high and low exponent, in relation with age and different education levels for the bilateral parietal regions; Figure S12: MMSE score changes according to age and education across different offset levels for the left hippocampus; Figure S13: MMSE score changes according to age and education across different offset levels for the left temporal region; Figure S14: MMSE score changes according to age and education across different offset levels for the right occipital region; Figure S15: MMSE score changes according to age and education across different offset levels for the bilateral temporal regions; Figure S16: Comparisons and significant differences between MMSE scores of participants with high and low offset, in relation with age and different education levels for the left hippocampus; Figure S17: Comparisons and significant differences between MMSE scores of participants with high and low offset, in relation with age and different education levels for the left temporal region; Figure S18: Comparisons and significant differences between MMSE scores of participants with high and low offset, in relation with age and different education levels for the left occipital region; Figure S19: Comparisons and significant differences between MMSE scores of participants with high and low offset, in relation with age and different education levels for the bilateral parietal region.

## Acknowledgments

We thank Hernandez et al. (29) for providing the publicly available dataset.

